# DeepCIP: a multimodal deep learning method for the prediction of internal ribosome entry sites of circRNAs

**DOI:** 10.1101/2022.10.03.510726

**Authors:** Yuxuan Zhou, Jingcheng Wu, Shihao Yao, Yulian Xu, Wenbin Zhao, Yunguang Tong, Zhan Zhou

## Abstract

**Motivation:** Circular RNAs (circRNAs) have been found to have the potential to code proteins. Internal ribosome entry sites (IRESs) are key RNA regulatory elements for the translation of proteins by circRNAs through a cap-independent mechanism. IRES can be identified by bicistronic assay, but the method is time-consuming and laborious. Therefore, it is important to develop computational methods for facilitating IRES identification, evaluation, and design in circRNAs.

**Results:** In this study, we proposed DeepCIP, a multimodal deep learning approach for circRNA IRES prediction, by exploiting both sequence and structure information. As far as we know, DeepCIP is the first predictor for circRNA IRESs, which consists of an RNA processing module, an S-LSTM module, a GCN module, a feature fusion module, and an ensemble module. The comparative studies show that DeepCIP outperforms other comparative methods and justify the effectiveness of the sequence model and structure model of DeepCIP for extracting features. We found that the integration of structural information on the basis of sequence information effectively improves predictive performance. For the real circRNA IRES prediction, DeepCIP also outperforms other methods. DeepCIP may facilitate the study of the coding potential of circRNAs as well as the design of circRNA drugs. DeepCIP as a standalone program is freely available at https://github.org/zjupgx/DeepCIP.

## Introduction

With the help of deep sequencing and computational analysis, researchers have found that circular RNAs (circRNAs) are a large class of RNAs with multiple functions, not only in animals and plants but also in viruses (Memczak *et al*., 2013; W. Zhao *et al*., 2019; J. Zhao *et al*., 2019). Although circRNAs are usually regarded as a class of noncoding RNAs that have a ring structure with covalent bonds without 5’ caps and 3’ poly(A) tails, increasing evidence indicates that circRNAs have the function of protein coding (Yang *et al*., 2018; M. Zhang, Huang, *et al*., 2018; Legnini *et al*., 2017; Xia *et al*., 2019). CircRNA-encoded proteins might be prevalent in several diseases, particularly exerting anti-tumor or tumor-promoting effects in human cancer (He *et al*., 2021; Pan *et al*., 2020; Jiang *et al*., 2021). This finding demonstrates the potential development and clinical application of the translation of circRNAs. Moreover, compared to linear mRNA, circRNA has better thermal stability, longer efficacy time, and more tissue and developmental-specific expression and is considered an ideal platform for the development of next-generation mRNA drugs (Sharma *et al*., 2021; Liu *et al*., 2022). Wei group (Qu *et al*., 2022) first established a technical platform for the efficient preparation of high-purity circRNAs in vitro and designed a circRNA vaccine encoding the spike protein receptor-binding domain (RBD) for SARS-CoV-2 and its variants. It is also believed that circRNAs also have a broad application in the prevention or treatment of infectious diseases, autoimmune diseases, and cancer.

The translation of circRNAs requires internal ribosome entry sites (IRESs), which is an RNA cis-acting regulatory element that can recruit small ribosomal subunits to the translation initiation site without the 5’ cap (M. Zhang, Zhao, *et al*., 2018; Pamudurti *et al*., 2017). IRES was first discovered in RNA virus genomes, such as those of poliovirus (PV) and encephalomyocarditis virus (EMCV) (Pelletier and Sonenberg, 1988; Jang *et al*., 1988). With the help of bicistronic assays, IRES has been widely discovered in both viral and cellular mRNA (Komar and Hatzoglou, 2005). Compared to cap-dependent translation, the mechanisms of IRES-mediated translation are relatively unknown. However, researchers believe that the primary sequence and RNA structure are functionally important for IRES activity, according to direct recruitment of the ribosome by the structured RNA, or partially indirect interaction with ribosome with the assistance of canonical initiation factors, as well as additional IRES trans-acting factors (ITAFs) (Stoneley and Willis, 2004; Komar and Hatzoglou, 2015). Moreover, compared with the linear RNA IRESs, circRNA IRESs contain higher GC content, lower minimum free energy (MFE), and are more structured in general (Chen *et al*., 2021). The different characteristics lead to different IRES activity in the linear RNA system and the circular RNA system.

IRESs are thought to play a significant role in a variety of cellular functions as well as a number of diseases. It has been estimated that about 10% of mRNAs may initiate translation by the IRES-mediated cap-independent mechanism (Stoneley and Willis, 2004; Komar and Hatzoglou, 2015). However, the research of IRES common features and functions has been hindered by the time-consuming and laborious traditional experimental procedures used to identify IRES elements. As a result, there are still very few IRESs that have been confirmed. Data-driven computational methods such as machine learning are increasingly being applied in biological data. Despite the fact that there are no universally conserved features in all IRESs, bioinformatics experts have still produced a number of prediction tools. A support vector machine (SVM)-based predictor named IRESPred was created by Kolekar et al.(Kolekar *et al*., 2016), which used 35 features for viral and cellular IRES prediction. However, the negative samples of the training dataset used in IRESPred are not the experimentally validated IRES-negative sequences. In 2016, a high-throughput bicistronic assay was designed (Weingarten-Gabbay *et al*., 2016) and identified thousands of novel viral and human IRES sequences. Machine learning techniques can be better used for IRES recognition, profit from the vastly increased amount of novel IRES sequences. IRESpredictor, a stochastic gradient boosting random forest regression model, was developed based on the high-throughput assay dataset to predict IRES activity using 6120 global and local sequence k-mer characteristics (Gritsenko *et al*., 2017). However, a lot of features could result in model overfitting and slow training times. Later, IRESfinder was introduced by Zhao *et al*. (2018), a logit model with carefully chosen 19 k-mer sequence characteristics that predicted only cellular IRESs using the human IRESs from the dataset. IRESpy, the most recent IRES prediction tool with greater performance and shorter training time, was developed based on 340 global k-mer sequence characteristics using the high-throughput assay dataset (Wang and Gribskov, 2019). The study in IRESpy also shows that the model based on sequence/structure hybrid features gets a slight performance increase over the sequence-based model. However, previous studies have been limited to the use of hand-crafted features that are hardly optimal, the role of structure in IRES prediction also needs to be further explored. Compared with the traditional machine learning algorithm, the deep neural network framework is under-explored but might be promising in IRES prediction. In addition, current IRES prediction methods are designed for linear mRNA, it’s still urgently needed to develop circRNA IRES prediction methods because of the differences in IRES activity in linear RNA and circRNA.

To address the limitations, in this study, we propose a novel multimodal deep learning-based model, called DeepCIP, which combines sequence and structural features to predict human and viral circRNA IRESs. To capture sequence information, we adopted Sentence-State LSTM (S-LSTM) to learn the high-latent sequence features automatically; on the other hand, we constructed the weighted RNA graph and used Graph Convolutional Network (GCN) to extract the secondary structure information from the graph of RNA. Through the feature fusion module, the features from sequence and structure were integrated to identify the circRNA IRES. To prove the effectiveness of DeepCIP, we benchmark DeepCIP and XGBoost model on the defined independent test set. The benchmarking results show the superior performance for DeepCIP over other comparative methods for circRNA IRES prediction.

## Materials and Methods

### Datasets

#### Data source

Weingarten-Gabbay *et al*. (2016) devised a high-throughput bicistronic assay that identified thousands of sequences with IRES activity from 55,000 oligos. On this basis, Chen *et al*. (2021) identified 17,201 eGFP(+) oligos and 23,654 eGFP(-) oligos by constructing an oligo-split-eGFP circRNA reporter construct according to the eGFP expression of oligo whether greater than background eGFP expression. Of these, 1,639 oligonucleotides (linearIRES) were captured as having specific IRES activity in the linear screening system and 4,582 oligonucleotides (circIRES) in the circular screening system. As a high-quality dataset is critical to the predictive performance of the model, filtering out higher confidence data for training the model is an essential step. We first set circIRES as the positive sample, and negative samples were obtained from eGFP(-) samples after exclusion of linearIRES and sequences with IRES activity greater than background activity (excluding those sequences with promoter activity of ⩾0.2 and splicing activity of ⩽-2.5) in the Weingarten-Gabbay *et al*. dataset. A total of 24,525 RNA sequences with 4,582 positive samples and 19,943 negative samples were obtained. Since these data contain a portion of synthetic sequences introduced to test the effect of specific mutations on IRES activity, we followed the approach of Weingarten-Gabbay *et al*. to retain from the dataset the native sequences tagged as “CDS screen”, “Human 5’UTR Screen”, “Viral 5’UTR Screen”, “Genome Wide Screen Elements”, “High Priority Genes Blocks”, “High Priority Viruses Blocks”, “rRNA Matching 5’UTRs”, and “IRESite blocks”. Finally, we obtained a dataset with 4,531 positive samples and 9,616 negative samples. Since the oligonucleotide library used for the IRES activity assay was artificially constructed, the length of all RNA sequences was 174 nt.

#### Training and test set split

Because of the lack of independent test datasets for circRNA IRES, we divided the dataset into a training dataset and a test dataset for training and evaluation of the model. We first collected human circRNA sequences from CircAtlas (Wu *et al*., 2020) and viral circRNA from Viruscircbase (Cai *et al*., 2021) and then mapped the RNA sequences in the dataset to the circRNA in the database by using blastn (Camacho *et al*., 2009). Finally, a total of 582 positive samples with 100% identity were identified. We then selected 582 negative samples randomly to form an independent test set together with the positive samples described previously for subsequent evaluation of the constructed model. The remaining 3,949 positive samples and 9,034 negative samples were used as the training set. The training and test datasets used in this study are shown in Table 1. To address the problem that the model is biased toward the class with more samples due to the imbalance of positive and negative samples, which reduces the generalization ability of the model, we used an approach that combines downsampling and model ensembles. Specifically, we randomly sampled the negative samples to obtain three subsets of the same number of negative samples as the positive samples, provided that the data were not wasted and that the data had no duplication where possible between the subsets. Afterward, all negative subsets were combined with the positive dataset to obtain three training subsets with 3,949 positive samples and 3,949 negative samples.

**Table 1.** Summary of the dataset used in this study.

| Dataset | CircIRES | NoncircIRES | Length (nt) |
| --- | --- | --- | --- |
| Training set | 3949 | 9034 | 174 |
| Test set | 582 | 582 | 174 |
| Total | 4531 | 9616 | 174 |

#### Ground-truth circRNA IRES dataset

We additionally gathered the experimentally verified circRNA IRES sequences with varying lengths from the literature as the positive samples composed a ground-truth dataset containing 10 IRES sequences from *Homo sapiens* circRNA and 4 IRES sequences from *Drosophila melanogaster* circRNA. The detailed information of the ground-truth data is shown in Table 2.

**Table 2.** The information of ground truth data.

| CircRNA name | Organism | Length | Reference |
| --- | --- | --- | --- |
| Circ-FBXW7 | *Homo sapiens* | 62 | Yang et al.(Yang *et al.*, 2018) |
| Circ-SHPRH | *Homo sapiens* | 127 | Zhang et al.(M. Zhang, Huang, *et al.*, 2018) |
| CircZNF609 | *Homo sapiens* | 211 | Legnini et al.(Legnini *et al.*, 2017) |
| Circβ-catenin | *Homo sapiens* | 107 | Liang et al.(Liang *et al.*, 2019) |
| CircPINTexon2 | *Homo sapiens* | 231 | Zhang et al.(M. Zhang, Zhao, *et al.*, 2018) |
| Circ-AKT3 | *Homo sapiens* | 216 | Xia et al.(Xia *et al.*, 2019) |
| CircFNDC3B | *Homo sapiens* | 77 | Pan et al.(Pan *et al.*, 2020) |
| CircMAPK1 | *Homo sapiens* | 54 | Jiang et al.(Jiang *et al.*, 2021) |
| Circ-HER2 | *Homo sapiens* | 149 | Li et al.(Li *et al.*, 2020) |
| CircSEMA4B | *Homo sapiens* | 130 | Wang et al.(Wang *et al.*, 2022) |
| CircMbl | *Drosophila melanogaster* | 456 | Pamudurti et al.(Pamudurti *et al.*, 2017) |
| CircPde8 | *Drosophila melanogaster* | 310 | Pamudurti et al.(Pamudurti *et al.*, 2017) |
| CircCdi | *Drosophila melanogaster* | 419 | Pamudurti et al.(Pamudurti *et al.*, 2017) |
| CircTai | *Drosophila melanogaster* | 568 | Pamudurti et al.(Pamudurti *et al.*, 2017) |

### RNA representation

#### RNA sequence processing

One-hot encoding is one of the most common ways of coding biological sequences. Here, for a given RNA sequence, we used one-hot encoding to represent the bases A, T/U, C and G as [1, 0, 0, 0], [0, 1, 0, 0], [0, 0, 1, 0] and [0, 0, 0, 1], respectively. Each sequence is denoted as a feature matrix of dimension [L × 4], where L denotes the length of the sequence and L = 174 in our dataset.

#### RNA secondary structure prediction and graph construction

Access to the structure of RNA is a difficult task because the structure of most RNA sequences is unknown. Some tools have been developed to predict the secondary and tertiary structure of RNA (Lorenz *et al*., 2011; Fu *et al*., 2022; Sato *et al*., 2021), where the ViennaRNA package is a classic code library containing a variety of programs for RNA secondary structure comparison and prediction. To obtain the RNA structure information, we used RNAplfold (Bernhart *et al*., 2006) in the ViennaRNA package (Version 2.5.1) to capture the dynamics of the RNA secondary structure. RNAplfold can calculate the locally stable secondary structure of the RNA by McCaskill‘s algorithm (McCaskill, 1990) and output the probability of RNA base-pairing. We set the parameters of RNAplfold as W=150, c=1e-3, and do not allow the structures with lonely pairs to produce (--noLP). Other parameters were used in the default settings.

The predicted base-pairing probabilities were used to construct the RNA-weighted graph G=(V, E, W). Each base is represented as a node V in the graph, where the features of each node are encoded using one-hot vectors. Edge E contains two different types of chemical bonding information: covalent bonds linking consecutive nucleotides along the RNA backbone and hydrogen bonds linking paired bases, where the weight W of the covalent bond is defined as 1 and the weight W of the hydrogen bond is equal to the probability of pairing the two bases to which it is attached.

### GNN for feature extraction

#### Sentence-State LSTM module

The model architecture used for RNA sequence feature extraction was Sentence-State LSTM (S-LSTM) (Y. Zhang *et al*., 2018). S-LSTM was proposed to address the limitations of BiLSTM utilizing the Graph LSTM. BiLSTM is a variant of recurrent neural networks, consists of a forward and a backward LSTM, and has been widely used in natural language processing. S-LSTM uses a gate mechanism similar to BiLSTM to control the information flow. They differ in that S-LSTM converts the text into a graph, treating each word as a word-level node w and adds a sentence-level node g to represent the complete sentence. Although S-LSTM acts on the sequence, the way the nodes aggregate and the message passing is similar to the operation of GNN. It was classified as a text GNN in Zhou *et al*. (2020)

At each time step t, information can be exchanged between the sentence-level node and each word-level node, and each word node will also exchange information with its context node. An S-LSTM state at each time step t can be defined as:

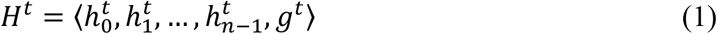

where 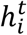is the substate of each word and *g*^*t*^ is the substate of the sentence.

As the time step t increases, each *h*_*i*_ captures an increasingly large n-gram context while exchanging information with g, making the contextual information learned by *h*_*i*_ and g increasingly rich. The final g can be used for the classification task. By default, each word node only exchanges information with the neighbor word node, which can be considered as using a window of size 1. However, increasing the window size can allow more information communication. Here, we set the window size to 3 and the time step to 9 by experiment.

The update process for the word state 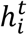 of S-LSTM in our model is given by the following equation:

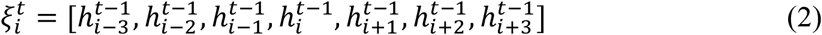

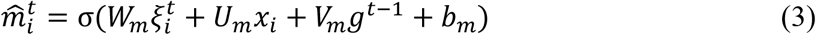

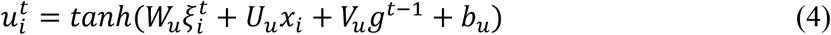

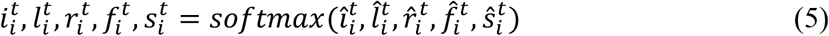

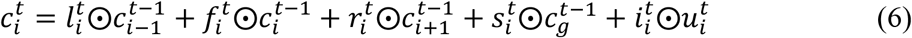

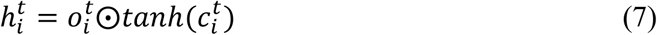

where 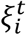is the concatenated vector of a context window, and 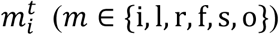 represents the different gates. 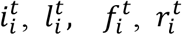, and 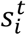 control information from the input *x*_*i*_, the left context cell 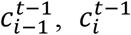, the right context 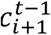, and the sentence context cell 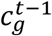, respectively; 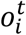 is the output gate. 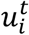is the actual input. *W, U, V*, and *b* are trainable parameters. *σ* is the sigmoid function.

The update process for the sentence state *g*^*t*^ of S-LSTM in our model is given by the following equation:

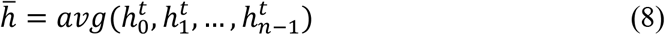

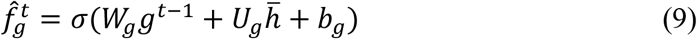

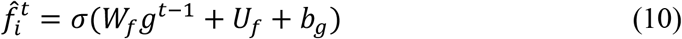

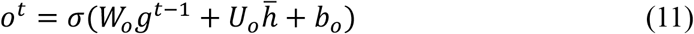

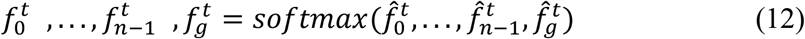

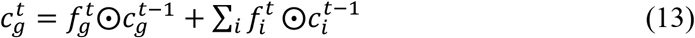

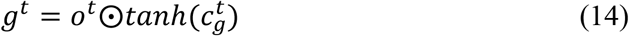

where 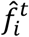and 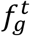 are the gates that are normalized to control the information from 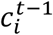 and 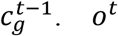 is the output gate. *W, U*, and *b* are trainable parameters.

#### GCN module

Graph neural networks (GNNs) are widely used in modeling the structure of molecules and proteins, but applications to RNA secondary structures are rare. We were inspired by Yan *et al*. (Yan *et al*., 2020) to extract secondary structure features using GNN. Each RNA secondary structure was represented as a weighted RNA graph. We make use of stacked multiple GCN layers to learn the feature vector of nucleotide nodes. To classify RNA sequences, aggregation of nucleotide node features in each RNA graph into graph-level embedding is needed. Global sum, max, and mean pooling are the most common strategies used to aggregate node features.

More specifically, the node features can be denoted as a matrix *X* ∈ *R*^*N*×*D*^, and the connectivity between nodes can be represented as an adjacency matrix *A* ∈ *R*^*N*×*N*^ with weight, where N is the number of nodes and D is the dimension of each node feature vector. The matrix X and A are the model inputs, and the GCN layerwise propagation rule as in Kipf & Welling (Kipf and Welling, 2017) is given by:

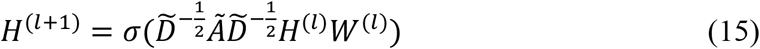

where *Ã* = *A* + *I*_*N*_, and *I*_*N*_ is the identity matrix. 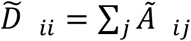is the diagonal degree matrix of *Ã. H*^(*l*)^ is the activation matrix, and *W*^(*l*)^ is the trainable matrix in the *l*^*th*^ layer. *σ*(·) is a nonlinear activation function.

Therefore, after several GCN layers, matrix X is converted to *Z* ∈ ℝ^*N*×*F*^ (F is the number of filters), and the formula of each node 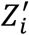 is given by:

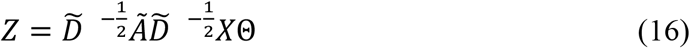

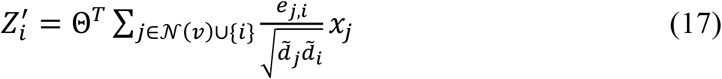

Here, Θ denotes a matrix of filter parameters, 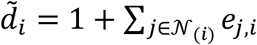, where *e*_*i,j*_ is the edge weight from node j to node i.

We choose to stack three GCN layers to learn the node-level embedding because too few layers will result in a small receptive field and too many layers may lead to over-smoothing. A global sum pooling layer is added after the final GCN layer to extract graph-level representations for the classification task because it obtained relatively better performance in our experiments. The specific experimental results are presented in “The structural model of DeepCIP” section.

### Feature fusion and model ensemble

To combine the sequence features and structure features derived from the S-LSTM and GCN modules, we concatenated them in the last dimension. Then, a classifier with a sigmoid function is used to output the predicted probabilities.

All training subsets were used separately for training, and three models were obtained. We apply a soft voting strategy to ensemble these three models. The voting mechanism is the most typical of the model ensemble methods, and the basic idea is to output the class with the most votes of all the classifiers. The voting method can be classified into two types denpending on the mechanism, where hard voting is the classifier directly gives the predicted label and soft voting is gives the predicted probability of the output label. We used soft voting to calculate the weighted sum of the probability of the three models, and the prediction label was then determined to be 0 or 1 based on a default threshold of 0.5. Here, we set the weights of the three models as equal.

### Nested cross-validation

Errica *et al*. (Errica *et al*., 2020) used nested cross-validation (CV) to fairly compare different GNN models. Briefly, a nested CV contains an external and an internal CV, where CV can choose the k-fold or holdout technique. K-fold CV denotes the random division of the dataset into *k*_*out*_ nonoverlapping subsets, where each subset takes turns as the test set, and the remaining subsets are used for training. Holdout CV means directly dividing the dataset into two mutually exclusive subsets, one for training and one for validation. Here, we apply a nested CV approach for hyperparameter tuning and model selection similar to Errica et al. The following steps are all performed on each of the three training subsets, which are described in the “Training and test set split” section. We use the k-fold technique with *k*_*out*_ = 5 for the external CV and use the holdout technique with a 90% training split and a 10% validation split in the internal CV. Specifically, we train each outer training fold, holding out a random 10% portion of the data as the validation set for executing the early stopping when there is no performance improvement after n epochs, and then test in the test fold. The final hyperparameter evaluation score is obtained from the mean of all test fold scores.

After completing the hyperparameter selection, we retrained and validated the model on all data and finally evaluated the model performance on the independent test set we constructed.

### Baseline methods

To assess the effectiveness of the proposed model and its various submodules, we compare it with other baseline methods using nested CV. The introduction of baseline methods is as follows:

#### TextCNN for sequence

TextCNN is a convolutional neural network for text classification, which is composed of an embedding layer, a convolution layer, a max-pooling layer, and a fully connected layer. where sequences are encoded as the one-hot vectors in the embedding layer, and we set n_filters=64 and filter_size=[2,3,4] in the convolution layer.

#### BiLSTM

BiLSTM is used to learn effective features from sequences, and it can be stacked in multiple layers. The single-layer, two-layer, and three-layer BiLSTMs are used for model comparison, where the hidden_size is set to 64.

#### TextCNN + annotate secondary structure

Here, RNAfold (Lorenz *et al*., 2011) is used to predict the RNA secondary structure and output a dot-bracket sequence representing the base pairing profile. bpRNA (Danaee *et al*., 2018) annotates the predicted RNA secondary structure, which resolves the base pairing information output by RNAfold into a detailed structure, providing relevant contextual annotation information including stems (S), hairpin loops (H), multiple loops (M), internal loops (I), bulges (B), and ends (E). This allows the RNA secondary structure to be represented as a sequence and fed into the TextCNN model for training. The setup of the TextCNN is described in the previous part.

#### Multilayer Perceptron (MLP) for the graph

This method applies a three-layer MLP with ReLU activations on the node features of the RNA graph, followed by a global sum pooling layer to learn the graph-level embedding. It differs from the GCN module in that it does not use relationships between nodes when learning node features and is a graph topology-independent model.

#### XGBoost

This model was used to develop the IRESpy tool (Wang and Gribskov, 2019). Here, 340 sequence k-mer (1-mer, 2-mer, 3-mer, and 4-mer) features are used to train the XGBoost model and set the same XGBoost hyperparameters as IRESpy. The parameter of scale_pos_weight is provided in the XGBoost model and is useful for unbalanced classes by controlling the balance of positive and negative weights. We trained two XGBoost models using the circRNA IRES training dataset, called XGBoost_weight for circ and XGBoost for circ, setting the parameter of scale_pos_weight to 3 and 1 (the same as IRESpy), respectively. We used 10-fold CV with the early stopping method to obtain the best num_boost_round parameter, and the best model was selected for comparison with our final model.

### Evaluation metrics

To evaluate the predictive performance of the model we proposed, we used five evaluation metrics similar to previous studies, including Accuracy (Acc), Sensitivity (Sn), Specificity (Sp), Precision, and Matthews correlation coefficient (MCC). The metrics are defined as follows:

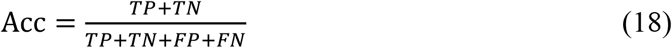

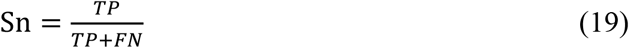

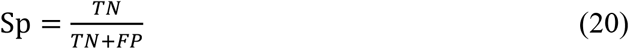

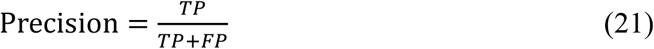

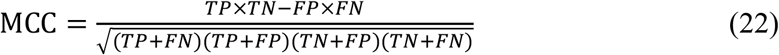

where TP, TN, FP, and FN denote the number of true positives, true negatives, false positives, and false negatives, respectively.

Moreover, we used the ROC (receiver operating characteristic) curve and PR (precision-recall) curve to intuitively evaluate the overall performance of our classification model.

## Results and Discussion

### The architecture of DeepCIP

We proposed DeepCIP, a multimodal deep learning method that is based on GNN, to extract features of RNA sequences and structures, and combine both features for circRNA IRES prediction. The architecture of DeepCIP is shown as Figure 1. Specifically, in the Ensemble Module, DeepCIP adopts the soft voting strategy to ensemble three fusion models trained by different datasets. Each fusion model comprises four modules, including the RNA processing module, S-LSTM module, GCN module, and feature fusion module. First, the RNA processing module is used to preprocess the input RNA sequence for sequence encoding, structure prediction, and RNA graph construction. Second, the S-LSTM module and GCN module are used to extract the features of the RNA sequence and RNA secondary structure, respectively. Finally, in the Features fusion Module, the features extracted from the RNA sequence and structure are fused. The fused feature passed through the fully connected layer and a sigmoid function, ultimately outputting a probability representing the likelihood that the input RNA sequence is a circRNA IRES.

**Figure 1.**
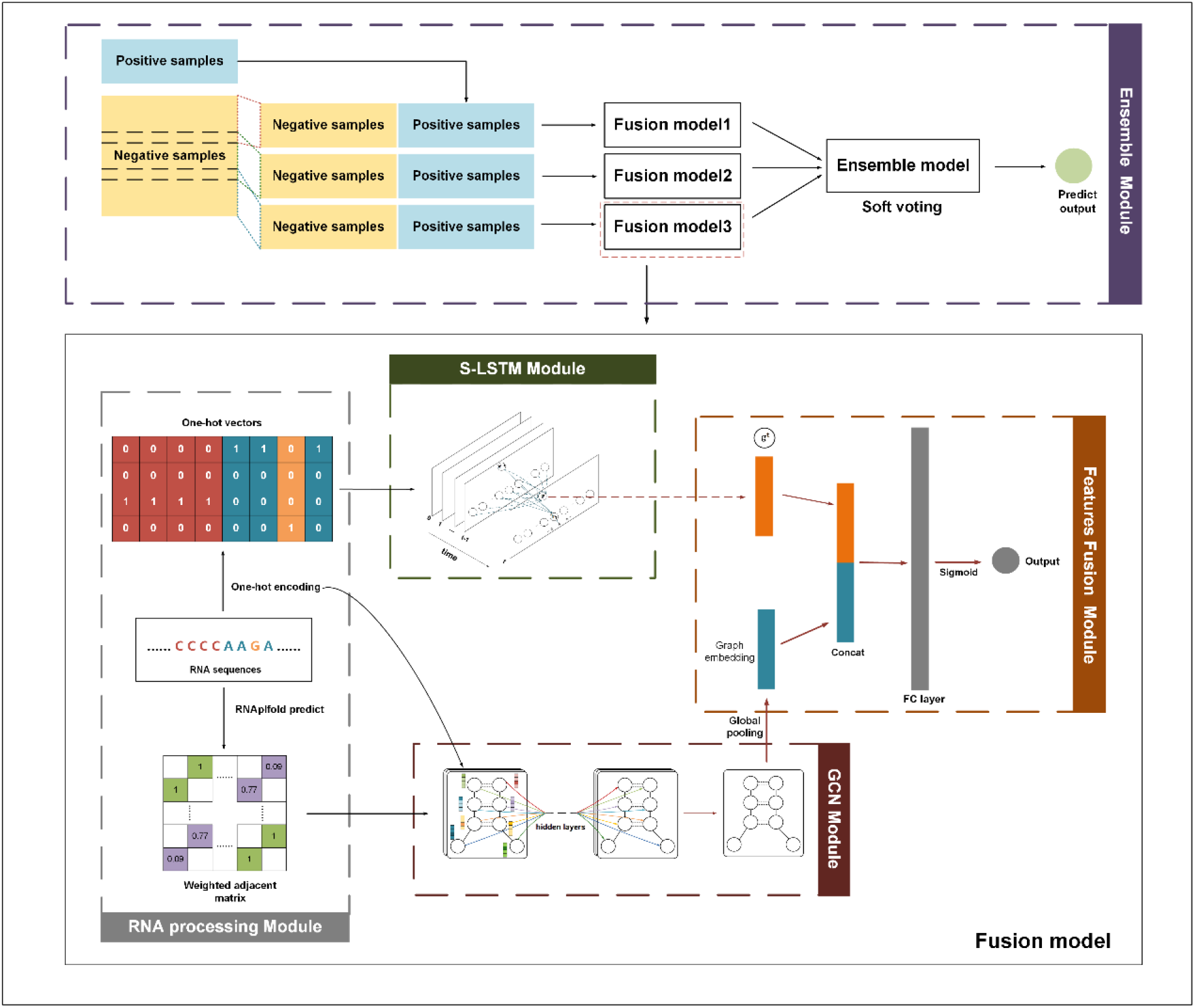
The architecture of DeepCIP method, including sequence processing, RNA graph construction, feature extraction, and model ensemble.

### The sequence model of DeepCIP

S-LSTM learns the representation of sequence by the Graph LSTM method. The performance of the sequence model is affected by many hyperparameters such as the number of sentence-level nodes, the window size, the time step, and the hidden layer size in the S-LSTM module. Here, we focus on the impact of window size and time step in the S-LSTM model, where window size ranged from {1, 2, 3}, and time step ranged from {5, 7, 9}. From the results of various S-LSTM settings depicted in Figure 2A, the setting of window size to 3 and the time step to 9 achieves the best value of average AUC. We further compared the performance of S-LSTM with TextCNN and BiLSTM under different numbers of stacked layers. Results indicate that S-LSTM outperforms other models in the value of average AUC (Figure 2B), which suggests that S-LSTM can represent circRNA IRES sequences more efficiently.

**Figure 2.**
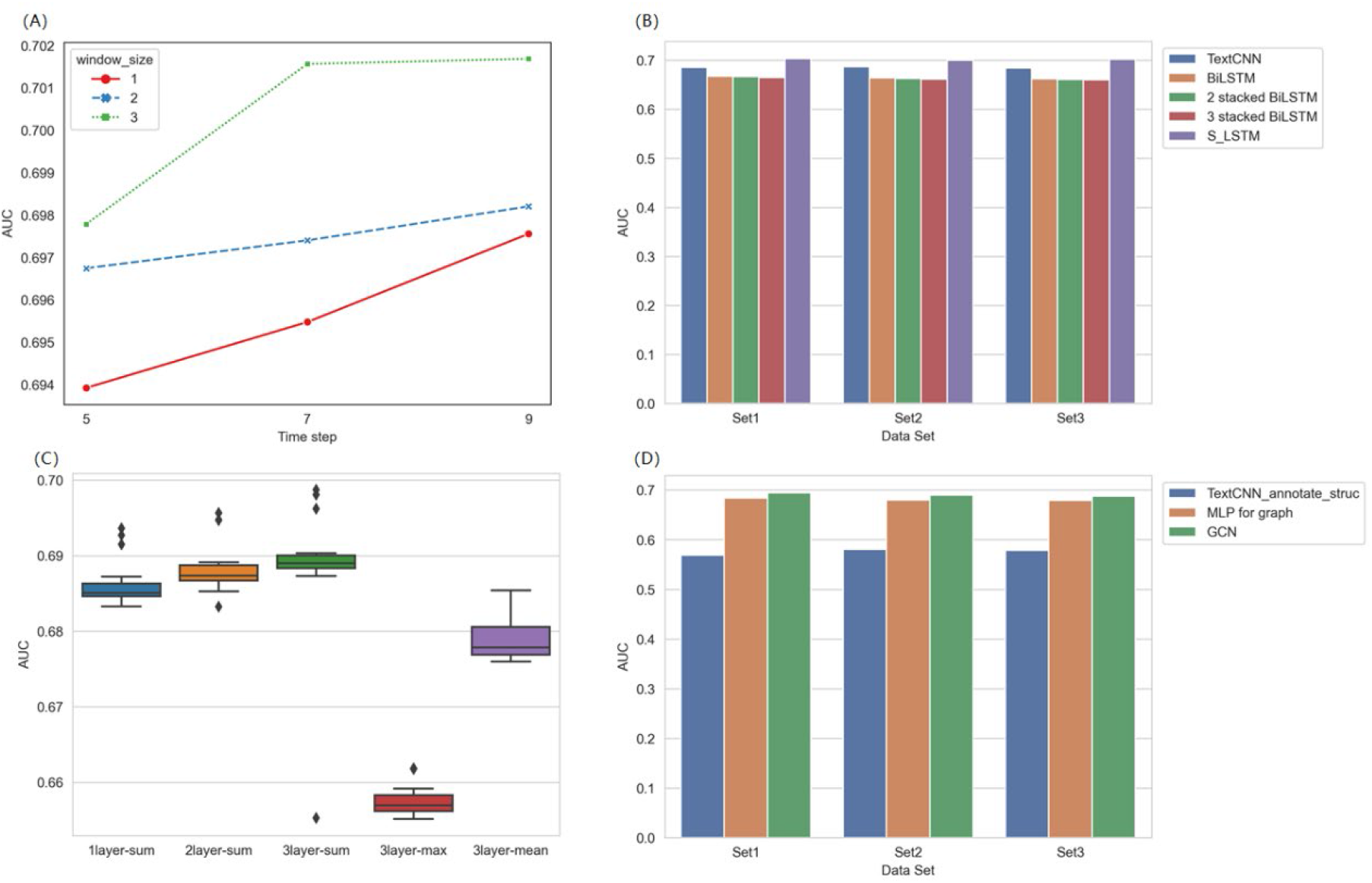
Hyperparameter optimization and model comparison of the sequence model and structure model. (A) Hyperparameter optimization of the sequence model. (B) Model comparison of the sequence model. (C) Hyperparameter optimization of the structure model. (D) Model comparison of the sequence model.

### The structural model of DeepCIP

The number of GCN layers and the readout function are crucial for feature extraction from the graph. Here, we construct a GCN module by varying the number of GCN layers and selecting different readout functions to study the impact of different configurations on model performance. The number of GCN layers ranged from {1, 2, 3}, and the readout strategy was chosen from {global sum pool, global max pool, global mean pool}. Results indicate that three layers of GCN followed by a global sum pool layer obtain the best performance concerning the AUC score (Figure 2C).

To investigate whether using the weighted RNA graph in our model could improve the representation ability circRNA IRES structure feature, we compared the performance of the weighted RNA graph + GCN, TextCNN + annotate secondary structure (TextCNN_annotate_struc), and MLP for graph. The comparative results of structure-based models show that the GCN module achieves the best performance (Figure 2D), which indicates that the weighted RNA graph can better represent the RNA secondary structure than the sequential annotation structure (GCN performs better than TextCNN_annotate_struc), and using different chemical bonding relationships between nucleotides is a better choice (GCN performs better than MLP for graph).

### Feature fusion and soft voting mechanism improve the performance of DeepCIP

To assess the importance of RNA structures in circRNA IRES recognition, the fusion model was compared with the model based on sequence features only on the independent test set. As shown in Figure 3A, the performance increases in the AUROC, AUPRC, ACC, SN, Precision, and MCC indicate the structural feature importance in circRNA IRES prediction.

**Figure 3.**
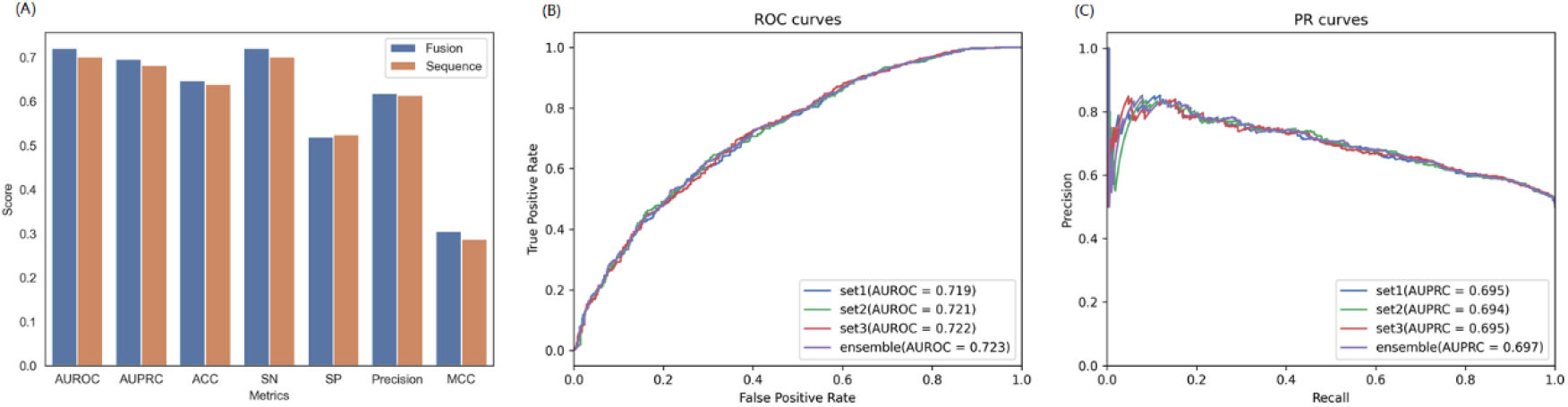
(A) The performance comparison between the models based on the sequence and structure fusion feature and based on the sequence feature only. (B) ROC curves of the ensemble model and single model. (C) PR curves of the ensemble model and single model.

Next, we performed tests on the independent test dataset to verify whether the soft voting mechanism can improve the predicted performance. Here, the soft voting method was used for three ensemble models trained by three training subsets. The comparative results between the three single models with the ensemble model indicate that the ensemble model has the best performance with AUROC of 0.723 and AUPRC of 0.697 (Figure 3B and 3C). However, the performance improvement is limited, which might be attributed to few models used for ensemble, and partial overlap between training subsets.

### Performance comparison between DeepCIP and XGBoost model

To evaluate the prediction performance of our model, we first carried out comparison experiments between our model and the XGBoost model with an independent test dataset. The comparative results are shown in Table 3. We can observe that our model gives higher performance than the XGBoost_weight model in terms of all metrics, achieving an AUC of 0.723, ACC of 0.646, SP of 0.512, SN of 0.780, Precision of 0.615, and MCC of 0.303. The reasons for this may be, on the one hand, that our model better represents the RNA structure and, on the other hand, that the features automatically extracted by deep learning have better representation ability compared to handcrafted k-mer features, which suggests that deep learning may be a beneficial choice to address the current situation where the common features of IRES are not yet fully clear.

**Table 3.** The comparative results of DeepCIP and XGBoost_weight for circRNA IRES prediction.

| Model | AUC | ACC | SP | SN | Precision | MCC |
| --- | --- | --- | --- | --- | --- | --- |
| XGBoost_weight for circ | 0.624 | 0.624 | 0.503 | 0.744 | 0.600 | 0.255 |
| Our model | **0.723** | **0.646** | **0.512** | **0.780** | **0.615** | **0.303** |

The boldface represents the best performance of different methods.

### Visualization of the predictive results of DeepCIP

To further investigate the effectiveness of the proposed DeepCIP, we explored the correlation between the predicted circRNA IRES probabilities and circRNA IRES experimental activities. The circRNA IRES activity is defined by the eGFP expression from Chen *et al*. experiment. In the independent test set, the activities of circRNA IRES are in the range of 0 to 7, where the background eGFP expression is 3.466387 (Chen *et al*., 2021). The result is illustrated in Figure 4A. We can observe that circRNA IRES with higher activity generally also have a greater prediction probability.

**Figure 4.**
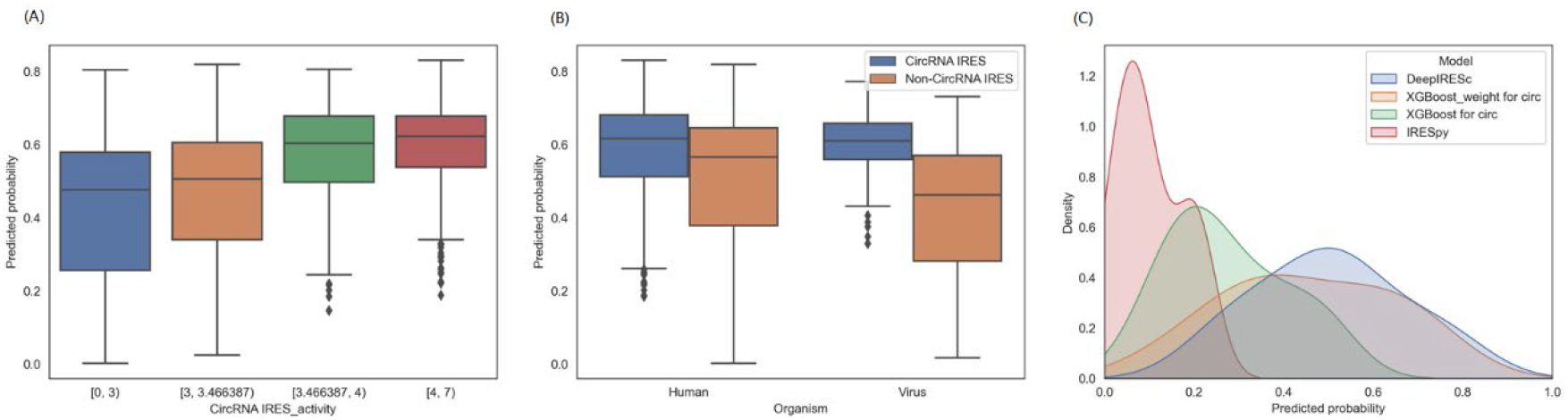
(A) The correlation between the DeepCIP predictive results and experimental results. (B) The predicted probabilities of DeepCIP on the human sequences and virus sequences from the independent test set. (C) The distribution of circRNA IRES prediction probability using different methods.

In addition, we visualized the predicted probabilities of DeepCIP on human sequences and virus sequences from the independent test set. As we can see in Figure 4B, the predicted probabilities of human and virus circRNA IRESs are both greater than noncircRNA IRESs in general. This demonstrates the ability of DeepCIP on both human and viral circRNA IRES predictions.

### Application to ground-truth circRNA IRES data

We further evaluated the predictive ability of our model on ground-truth data that are not included in the training dataset and compared the predicted results of DeepCIP with other models including XGBoost_weight for circ, XGBoost for circ, and IRESpy. The predicted results are shown in Table 4. For the results of our model, if the prediction threshold is set to 0.5, 6 out of 10 RNA sequences from *Homo sapiens* are predicted to be circRNA IRES, and 1 out of 4 RNA sequences from *Drosophila melanogaster* are predicted to be circRNA IRES, performs better than other methods. The poor performance of DeepCIP on RNA sequence from *Drosophila melanogaster* might be that there are differences in circRNA IRES characteristics among different species, and our training dataset contains only RNA sequences from humans and viruses. Moreover, Figure 4C shows the distribution of circRNA IRES prediction probability using different methods for the human circRNA IRESs from ground-truth data by a kernel density estimate (KDE) plot. We can observe that DeepCIP reaches a promising performance when compared to XGBoost-based models, and the prediction performance of IRESpy (used for linear RNA IRES identification) is lower than other methods used for circRNA IRES prediction in real circRNA IRES data, which indicates the importance to develop methods specifically for circRNA IRES identification.

**Table 4.** Prediction results of different models on the ground-truth dataset.

|  | DeepCIP | XGBoost_weight for circ | XGBoost for circ | IRESpy |
| --- | --- | --- | --- | --- |
| Circ-FBXW7 | **0.571** | **0.573** | 0.349 | 0.069 |
| Circ-SHPRH | 0.395 | 0.427 | 0.214 | 0.047 |
| CircZNF609 | **0.551** | **0.603** | 0.342 | 0.019 |
| Circβ-catenin | 0.432 | 0.454 | 0.236 | 0.207 |
| CircPINTexon2 | 0.264 | 0.342 | 0.197 | 0.200 |
| Circ-AKT3 | 0.281 | 0.280 | 0.137 | 0.027 |
| CircFNDC3B | **0.527** | 0.310 | 0.204 | 0.197 |
| CircMAPK1 | **0.501** | 0.147 | 0.113 | 0.111 |
| Circ-HER2 | **0.766** | **0.690** | 0.471 | 0.083 |
| CircSEMA4B | **0.727** | **0.711** | 0.486 | 0.074 |
| CircMbl | 0.009 | 0.123 | 0.056 | 0.067 |
| CircPde8 | 0.394 | 0.375 | 0.197 | 0.024 |
| CircCdi | 0.130 | 0.371 | 0.188 | 0.042 |
| CircTai | **0.597** | **0.546** | 0.341 | 0.029 |

The boldface represents the predictive probability greater than 0.5.

## Conclusions

In this study, we propose DeepCIP, a multimodal deep learning method based on the sequence and structure features for circRNA IRES prediction, which can identify the variable lengths of circRNA IRES sequences. Since the function of IRES relies on the RNA structure, we constructed a weighted RNA graph to model the RNA secondary structure and extracted structural features based on GCN. Meanwhile, we innovatively used S-LSTM to learn a global sentence-level node to characterize the entire RNA sequence, thus better modeling the contextual information for the classification task. The experimental results show that the S-LSTM module and GCN module achieve the best performance in extracting features from sequences and structures compared with a variety of deep learning methods. By integrating the sequence and structure information, DeepCIP can capture the features of circRNA IRES more effectively. Moreover, to solve the problem caused by imbalanced data, we applied negative data downsampling and soft voting strategies. DeepCIP is the first tool specifically developed for circRNA IRES prediction and is the first to use deep learning methods for the identification of IRES sequences. The experimental results show that DeepCIP can predict human and viral circRNA IRES with different lengths effectively. We suggest that this tool be applied to investigate the translational regulation of circRNAs and the subsequent design and application of circRNA based drugs, such as vaccines, cytokines, and so on.

This study still has several limitations. First, the data used for model training were fully designed RNA sequences, all 174 nt in length. However, the real circRNA IRES sequences have different lengths. Second, the mechanism of most IRESs is related to the recruitment of ribosomes, and regulation by eukaryotic initiation factors (eIFs) and ITAFs (King *et al*., 2010), considering only sequence and structure features, may be insufficient to interpret the mechanism of IRESs. Therefore, future work is needed to further consider the information on RNA-RNA and RNA-protein interactions. Furthermore, extension work to optimize the nucleotide encoding method and model architecture should be performed in further studies.

## Data availability

The software are accessible at https://github.org/zjupgx/DeepCIP. The raw experimental data applied in this study are available in Gene Expression Omnibus (https://www.ncbi.nlm.nih.gov/geo/), and can be accessed with GSE178718.

## Acknowledgments

We thank the Information Technology Center and State Key Lab of CAD&CG, and the Innovation Institute for Artificial Intelligence in Medicine, Zhejiang University for the support of computing resources.

## Funding

This work was supported by the R&D Program of Zhejiang University Innovation Institute for Artificial Intelligence in Medicine - Aoming (Hangzhou) Biomedical Co., Ltd. Joint Laboratory [Grant No. 20220210], the R&D Program of China Jiliang University - Aoming (Hangzhou) Biomedical Co., Ltd. Joint Laboratory [Grant No. 20211008] and the R&D Contract of Aoming (Hangzhou) Biomedical Co., Ltd [Grant No. 2021-05].

## Competing interests

Y. Tong is the founder of Aoming (Hangzhou) Biomedical Co., Ltd. All other authors declare no competing financial interests.

## Notes

### Competing Interest Statement

Y. Tong is the funder of Aoming (Hangzhou) Biomedical Co., Ltd. All other authors declare no competing financial interests.

### Summary of Updates

There is a mistake in the author name for Yunguang Tong. We have modified it.

